# Mixed matrix factorization: a novel algorithm for the extraction of kinematic-muscular synergies

**DOI:** 10.1101/2021.08.05.455189

**Authors:** Alessandro Scano, Robert Mihai Mira, Andrea d’Avella

## Abstract

Synergistic models have been employed to investigate motor coordination separately in the muscular and kinematic domains. However, the relationship between muscle synergies, constrained to be non-negative, and kinematic synergies, whose elements can be positive and negative, has received limited attention. Existing algorithms for extracting synergies from combined kinematic and muscular data either do not enforce non-negativity constraints or separate non-negative variables into positive and negative components. We propose a mixed matrix factorization (MMF) algorithm based on a gradient descent update rule which overcomes these limitations. It allows to directly assess the relationship between kinematic and muscle activity variables, by enforcing the non-negativity constrain on a subset of variables. We validated the algorithm on simulated kinematic-muscular data generated from known spatial synergies and temporal coefficients, by evaluating the similarity between extracted and ground truth synergies and temporal coefficients when the data are corrupted by different noise levels. We also compared the performance of MMF to that of non-negative matrix factorization applied to separate positive and negative components (NMFpn). Finally, we factorized kinematic and EMG data collected during upper-limb movements to demonstrate the potential of the algorithm. MMF achieved almost perfect reconstruction on noiseless simulated data. It performed better than NMFpn in recovering the correct spatial synergies and temporal coefficients with noisy simulated data. It also allowed to correctly select the original number of ground truth synergies. We showed meaningful applicability to real data; MMF can also be applied to any multivariate data that contains both non-negative and unconstrained variables.

## 1. Introduction

In recent years, several motor control studies have focused on the evaluation of muscle or kinematic coordination patterns. They have often been based on the premise that the Central Nervous System (CNS) relies on a limited number of modules [1], possibly implemented at neural level [2], to simplify the production of movement. Consequently, by properly recruiting spatial modules scaled by temporal activation coefficients, the CNS would exploit a reduced set of pre-shaped neural pathways, called synergies, to achieve a large variety of motor commands. This view implicitly assumes that, if synergies are encoded at neural level, a unique set should be used across a variety of movements or, at least, task-specific sets should underlie movements requiring similar motor commands. Thus, different movements within a task (e.g., reaching movements in different directions – [3]) may be generated by one set of synergies, but different tasks may require different or additional synergies.

The notion of motor modules has been widely exploited following two main lines that have remained so far mostly separated: muscle synergies and kinematic synergies. While the idea of a modular organization of the motor system has deep roots in neurophysiology [4] [5] [6], two decades ago Bizzi and collaborators introduced a simple yet quantitative model for the generation of muscle patterns through the combination of muscle synergies [1], [7], [8], [9], inspired by the organization of the spinal cord [10], [11], which has since become the standard approach to characterize muscle coordination patterns. The algorithm most commonly used for synergy extraction is non-negative matrix factorization (NMF) [12]. Under the assumption of non-negativity, which naturally applies to muscle activation signals, NMF provides a sparse representation of modular control elements by factorizing multiple muscle activation waveforms, in the most commonly used decomposition approach, as the product of non-negative, time-invariant muscle weighting vectors (spatial synergies) and non-negative time-varying coefficients. Additional approaches for synergy extraction include temporal [13], time-varying [3], [9] and the space-by-time [14] synergy models and algorithms.

In parallel, starting from the work of Santello and collaborators [15], a comparable line was followed to describe the synergistic recruitment of kinematic modules by grouping co-varying joint angles. We note that, differently from EMG, kinematic waveforms may be either positive or negative (modelling the flexion and extension of a joint). Kinematic synergies can be identified with algorithms such as principal component analysis (PCA). This approach was exploited by many authors, e.g. for locomotion [13] and hand grasps [16]. Depending on the body segment under analysis, one approach might be preferable in respect to the other (e.g., kinematic synergies seem to be more effective and appropriate to describe hand patterns rather than muscle synergies, that are instead popular for the analysis of the upper-limb and locomotion).

Although it may be easier to analyze the control of different body segments or gestures separately in one of the two domains, it is clear that muscular and kinematic analysis are related, yet complementary, approaches. However, despite the potential advantages of the combination of the two approaches, only a few studies have analyzed the relationship between muscular and kinematic variables in terms of motor modules. Tagliabue and collaborators [17], analyzing two-digit grasping, found that a reduced number of modules (2–3) was needed to explain the largest part of the variation for each grasp and suggested a correlation between muscle and kinematic primitives, justifying synergy-based analysis in both domains. Russo and collaborators [18] investigated the relationship between the dimensionality of joint torques and muscle patterns for reaching movements. However, muscular and kinematic variables have rarely been analyzed together in the framework of synergistic control. Even if the dynamic relationship between muscle activation and kinematic variables is in general non-linear, in a simple arm model it is possible to generate any end-effector trajectory thought linear combinations of muscle synergies [19]. Thus, it is reasonable to hypothesize that the identification of combined “kinematic-muscular synergies” may provide insights on the functional role of motor modules and on the relationship, even if approximated, between muscle synergies and the motor output at the kinematic or kinetic level. This relationship could possibly be captured by separately extracting kinematic and muscle synergies, but only qualitatively and indirectly. In this work, we propose a novel approach that allows to extract combined “kinematic-muscular synergies”.

Combining an EMG and kinematic-based analyses under a unique domain requires a new factorization algorithm. Using algorithms without non-negativity constraints such as PCA, factor analysis (FA), or independent component analysis (ICA) [20], [21] produces spatial synergies with negative weightings for the muscular variables that do not much the physiological non-negativity of muscle activation and cannot be easily interpreted. Moreover, PCA introduces constrains of orthogonality on extracted synergies that are also not physiologically justified. Using the NMF algorithms is also problematic. In fact, while EMG activation are non-negative, kinematic waveforms are by nature not subject to this constraint. In example, bell-shaped velocity profiles found in human articular motion are coupled with biphasic (positive-negative) acceleration waveforms; in other words, at articular level, each joint can be flexed or extended. A relevant attempt to include negative waveforms in combined muscular-kinetic synergies was made by Ting and collaborators [22], [23]. Positive and negative components of each force waveform were considered two separate non-negative waveforms, and “functional muscle synergies” [23], comprising both muscular and force signals, were extracted with NMF (from here, we will refer to this algorithm as NMFpn). However, NMFpn may not correctly identify functional muscle synergies, since data generated by non-negative combinations of unconstrained force vectors may require a larger number of synergies when positive and negative force components are considered as separate dimensions. To the best of our knowledge, there is currently no other algorithm capable of extracting synergies from combined non-negative and unconstrained (positive and negative) signals (e.g., kinematic-muscular).

To fill this gap, we propose a novel factorization algorithm to extract kinematic-muscular synergies, and, potentially, any synergy from combined data from multiple domains with and without non-negativity constraints. The innovative feature is that we do not constrain all the variables of a spatial synergies to be non-negative; on the contrary, there can be any mixture of constrained and unconstrained variables. Thus, when applied to combined muscular and kinematic data, we constrain EMG synergy components to be non-negative, while kinematic synergy components are left unconstrained. To do so, the algorithm substitutes the multiplicative update rule of the original NMF algorithm with a gradient descent update rule, with a regularization term to reduce redundancy and enforce sparseness on the extracted synergies. Our paper introduces the mathematical formulation of the mixed matrix factorization (MMF) algorithm, presents its validation through comprehensive simulations of data generated by non-negative combinations of known kinematic-muscular synergies (ground truth), investigates its robustness by assessing the effects of noise on the accuracy of the reconstruction of ground truth synergies and temporal coefficients, compares its performance with respect to the current state-of-the-art, and, finally, shows its applicability to real scenario of upper-limb movements.

## 2. Methods

### 2.1 Study design

The paper is organized as follows. First, we introduce the formulation of the MMF algorithm and we identify valid ranges for the parameters of the algorithm. Then, we present a comprehensive set of simulations to quantify the accuracy of the reconstruction of spatial and temporal features on biologically-inspired simulated data corrupted by different amounts noise, in order to validate the algorithm. We also provide comparison with the NMFpn algorithm [23] to show how our algorithm overcomes the limitations introduced by the separation of unconstrained components into distinct positive and negative components. Lastly, we apply the MMF algorithm to real data to demonstrate its applicability and potential for identifying kinematic-muscular synergies.

### 2.2 Mixed Matrix Factorization Algorithm

We developed and implemented a gradient descent algorithm for extracting synergies from a data matrix with a mixture of unconstrained and non-negative components. As the standard Non-Negative Matrix Factorization (NMF, [12]) algorithm commonly used for muscle synergy extraction, the new Mixed Matrix Factorization (MMF) algorithm factorizes a data matrix **X** (with time samples of several variables arranged along the *rows*). However, while NMF required all variables to be non-negative, MMF can be applied when only as subset of variables are non-negative, thus expanding its range of applications. In NMF, the original data matrix is decomposed with an iterative update procedure to estimate **C** (temporal coefficients matrix) and **W** (spatial synergies matrix), under the assumption that **X = WC +** ε, with ε residual noise. In the original NMF, **C** and **W** are both constrained to be non-negative. In the MMF algorithm, we consider vector data (**x**) with the first *k* components unconstrained (x_*i*_ ∈ **ℛ** for *i* = 1, …, *k*) and the remaining *m* components non-negative (x_*i*_ ∈ **ℛ**^+^ for *i* = *k* + 1, …, *k* + *m*). We want to decompose collections of such data (e.g. samples of kinematic and muscle data collected simultaneously) as combination of *n* synergies (w_*i*_ with w_*ij*_ ∈ **ℛ** for *j* = 1, …, *k* and w_*ij*_ ∈ **ℛ**^+^ for *j* = *k* + 1, …, *k* + *m*):

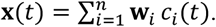

In matrix form (**X** = [**x**(1) … **x**(*S*)], **W** = [**w**_1_… **w**_*n*_], **C** = [**c**(1) … **c**(*S*)], with *S* number of samples):

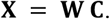

We decompose by minimizing the following cost function:

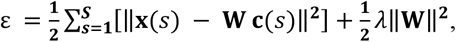

where the first term is the squared reconstruction error and the second a regularization term, enforcing minimum norm solutions. Taking the gradient of the error with respect to the components of **W** we obtain:

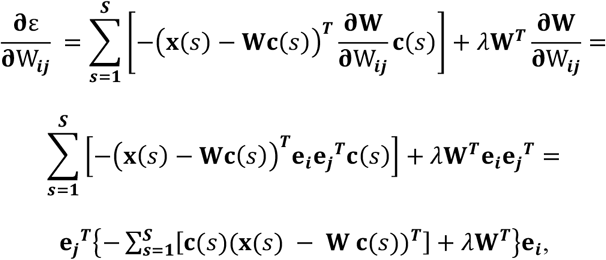

where **e**_*i*_ and **e**_*j*_ are unit vectors along the direction of the i^th^ component of **x** and the j^th^ components of **c** respectively. In matrix form:

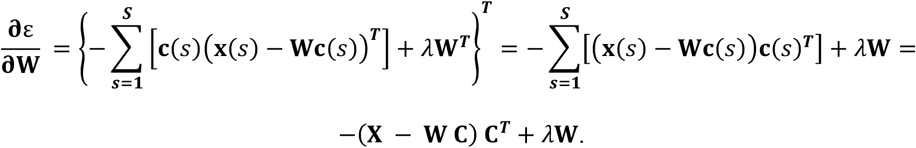

Similarly, for the gradient of the error with respect to the components of **C**:

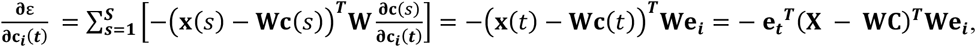

where **e**_*t*_ is a unit vector along the direction of the *t*^th^ sample (column of the data matrix). In matrix form:

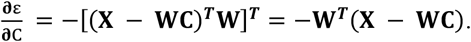

Then, the iterative update rules are (with learning rates *μ*_*W*_ and *μ*_*C*_):

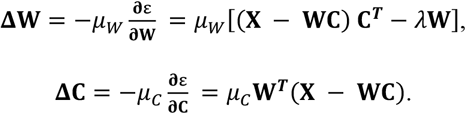

At the end of the extraction, **W** were normalized to unit vectors, and **C** were rescaled proportionally. The innovative feature of the MMF algorithm is that the gradient descent update rule allows a selected number of **W** variables to be either positive, zero or negative. For kinematic-muscular synergies, this allows to consider when a joint is flexed or extended.

The regularization term has been introduced to avoid non-physiological solutions. In fact, without such term, the algorithm might converge to solutions in which two or more synergies have components for the unconstrained variables with opposite signs and, simultaneously, overlapping temporal coefficients. In such solutions, the contribution of the components with opposite signs in different synergies may show destructive interference. Such cancellations may lead to synergies with large components for the unconstrained variables (e.g., the kinematic variables), which do not have a clear physiological interpretation. In contrast, the regularization term penalizes large synergy components even if, due to cancellations, they do not affect the reconstruction of the data, and therefore the reconstruction error, leading to sparser solutions for **W** and **C**. Moreover, such cancellations in the kinematic variables would be essentially associated to co-activations in muscles generating joint torques that cancel each other (i.e., co-contractions) and the regularization term thus has a physiological interpretation as effort associated to co-contraction. Increasing the relative contribution of the regularization term in the cost function, i.e., increasing the parameter *λ*, reduces or eliminates the occurrence of cancellations. However, when *λ* increases, the reconstruction quality, measured by R^2^ defined as the fraction of total variation explained by the synergy combinations, may decrease. The proper *λ* should then be chosen as a trade-off between sparseness and reconstruction accuracy.

We devised a procedure to tune the *μ*_*W*_, *μ*_*C*_, and *λ* parameters, as they concur to the magnitude of the update steps of the gradient descent. As the magnitude of ∥**X**∥, which depend on the dimension of the dataset, affects the magnitude of the gradient descent step, we expressed the parameters of the update rules as:

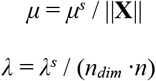

where *n*_*dim*_ = *m* + *k* is the number of rows of W (i.e., the total number of EMG and kinematic variables), and *n* is the order of the factorization (i.e., the number of extracted synergies). The scaled parameters *μ*^*s*^ and *λ*^*s*^ are advantageous to use because they are independent of the specific dimensions of the dataset, ensuring that a similar gradient descent step is used in different datasets. Notwithstanding this scaling, *μ*^*s*^ and *λ*^*s*^ are parameters that should be tuned depending on the dataset and on the desired level of sparseness of the solutions **W** and **C**.

The output of our novel algorithm are multi-domain kinematic-muscular spatial synergies and time-varying combination coefficients. Each extracted synergy is a vector that contains the components (or weights) for a mixture of the muscular and kinematic variables. Such vectors represent directions, in the combined muscular and kinematic space, in which there is covariation among the variables, i.e. a linear coordination across the two physical domains. The time-varying coefficients represent, for each repetition, the recruitment of each time-invariant synergy over time.

### 2.3 Non-Negative Matrix Factorization applied to separate positive and negative components

To show the effectiveness of our novel algorithm, in this study we have also implemented the approach used by Torres-Oviedo and collaborators [23] to extract functional muscle synergies from combined EMG and force data with the NMF algorithm. This approach (NMFpn) uses NMF on combined non-negative and unconstrained data after separating the negative and positive waveforms of the unconstrained signal and treating them as separate waveforms. The positive part of the signal (e.g. the positive part of each component of the force vector) is used as input for the NMF; the rectified negative components (e.g. the negative part of each component of the force vector) are used a separate input to the algorithm. In this study, the NMFpn approach was used to extract kinematic-muscular synergies using the standard NMF implementation based a multiplicative update rule [12]. To apply NMF to vector data (x) with the first *k* components unconstrained and the remaining *m* components non-negative, each unconstrained component is separated into a positive part 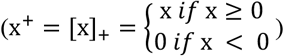 and a negative part 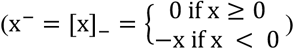, and NMF is applied to expanded *2k + m* dimensional vector data (**y**) with only non-negative components (y_*i*_ ∈ **ℛ**^+^, with y_*i*_ = [x_*i*_]_+_ *for i* = 1, …, *k*; y_*i*_ = x_*i*_ *for i* = *k* + 1, …, *k* + *m*; y_*k*+*m*+*i*_ = [x_*i*_]_−_ *for i* = 1, …, *k*). Both spatial synergies (**Z**) and temporal components (**C**) were initialized with small values. Then the update rules for spatial synergies and temporal components were iterated to minimize the chosen cost function until a termination condition was achieved. The Euclidean distance between the input and the reconstructed signal was used as cost function:

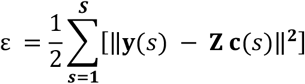

The multiplicative update rule for the synergies can be written in matrix form [12] as:

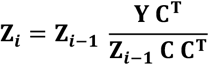

and the multiplicative update rule for the temporal components is:

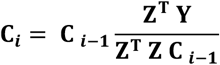

After the extraction of the synergies, the components of the synergies corresponding to the separated negative and positive components of the original data were recombined (w_*i*_ = z_*i*_ − z_*k*+*m*+*i*_ *for i* = 1, …, *k*; w_*i*_ = z_*i*_ *for i* = *k* + 1, …, *k* + *m*) into a synergy matrix (**W**) with the same number of rows as the dimension of the original data.

### 2.4 Validation on simulated data

We performed a comprehensive set of simulations to validate the algorithm, as shown in detail in Fig. 1.

**Figure 1:**
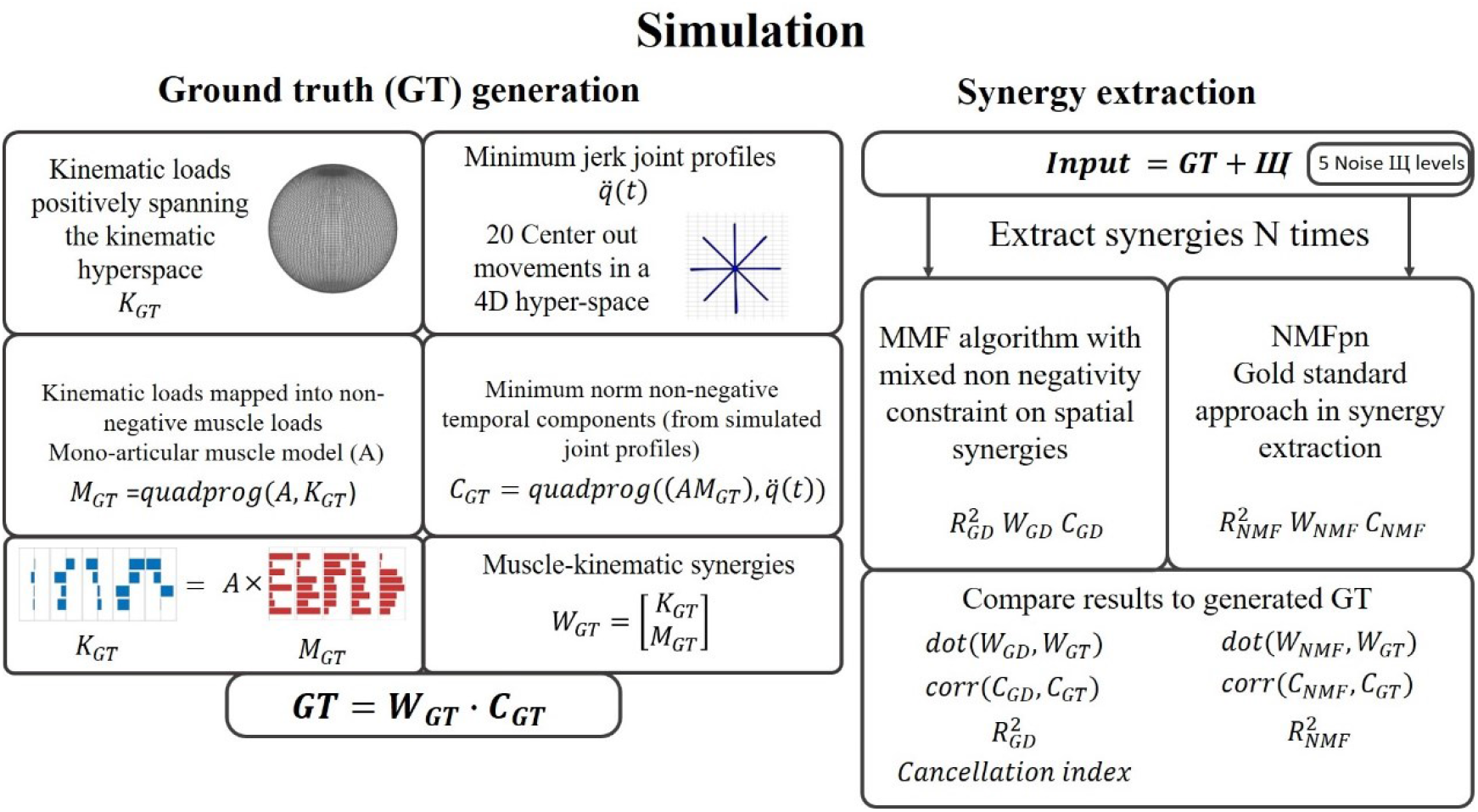
Simulation overview. Each simulation was divided into 3 parts. The first was the generation of a set of ground truth spatial synergies W_GT_ and combination coefficients (or temporal components) C_GT_. In the second part of the simulation, we extracted the synergies with the MMF algorithm and with the NMFpn algorithm from the simulated data W_GT_·C_GT_ corrupted by additive noise. Both algorithms included an initialization of the solutions (W_0_ and C_0_), iterative update based on gradient descent (MMF) or matrix multiplication (NMFpn), and a termination condition. Additionally, both algorithms were used to extract synergies with different noise levels to verify their robustness. Finally, the last step of the simulation was the comparison of the extracted synergies and coefficients with the ground truth ones. This assessment was based on comparing the reconstruction R^2^, the similarity between extracted and ground truth spatial synergies and combination coefficients, a cancellation index.

In particular, our main aim was to determine whether the algorithm recovered a known set of simulated synergies and combination coefficients used to generate the data. We also wanted test the effect of several noise levels added to the input data on the extracted synergies.

#### 2.4.1 Ground Truth (GT) generation

Our procedure began with the generation of a set of ground truth synergies (W_GT_) and combination coefficients (C_GT_) so that data input to the factorization algorithm (**X**) could be computed as:

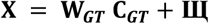

where **щ** is additive noise.

Using **X** as input, we extracted **W** and **C** in several conditions (W and C initializations, different noise levels, and number of iterations) and compared them to the original **W**_**GT**_ and **C**_**GT**_. So, our first step was to generate the ground truth **W**_**GT**_ and **C**_**GT**_. In previous related works (e.g. [20]), random synergies and coefficients were used as ground truth. However, it has been reported that biomechanical and task constraints affect the results of synergy extraction algorithms [24]. Thus, we generated data according to simple yet reasonable biomechanical constraints. At the same time, we deemed unnecessary to use a complex musculoskeletal model to validate our algorithm, and we chose a simple linear model. Indeed, our aim was to generate ground truth synergies and temporal coefficients with biologically inspired constraints, rather than proposing or validating any biomechanical model.

To generate synergies and combination coefficients coherent with the specific features of the algorithm (i.e., only a subset of non-negative variables) and biologically-inspired (i.e., reflecting the fact that movement kinematics is affected by muscle activation), we assumed a linear relationship between muscular components (**M**) and kinematic components (**K**) of the synergies. We defined a matrix **A** as the “mapping operator” that associates muscle activation vectors (**m**) to vectors of kinematic variables such as joints accelerations (**k**). Well aware of the biomechanical limitations of this model, **A** can be thought as “muscle moment arm matrix”.

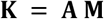

where **K** has dimension *k* × *n* (number of articular joints × number of synergies), **M** has dimension *m* × *n* (number of muscles × number of synergies), and **A** has dimension *k* × *m*.

Each **A** element could be either positive or negative (depending on the sign of the torque generated by each muscle around each joint), or zero (in case a muscle did not act on a specific joint). The elements of **A**, essentially expressing the pulling directions in joint space of each muscle) were generated simulating a set of monoarticular muscles pulling each joint in either flexion or extension directions. We considered a four-dimensional kinematic space and an eight-dimensional muscular space, with a pair of agonist-antagonist muscles for each joint:

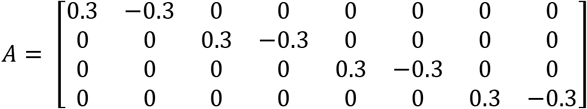

Then, we chose the kinematic components of the simulated synergies (**K**) as random directions in kinematic space, assuring that the set of directions included in **K** represents a basis would positively span the kinematic *k*-dimensional space [25]. The corresponding muscular components of the simulated synergies (**M**) were determined from **K = A M**. Since, in general, infinite solutions for **M** can be found, we selected non-negative solutions with minimum norm (with quadratic programming optimization, using Matlab function *quadprog*). For each kinematic component of the *i*-th synergy **k**_**i**_, we identified the corresponding muscular component **m**_**i**_ as the solution to the following minimization problem:

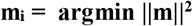

under the constraints **k**_**i**_ **= Am**, m_*ij*_ ≥ 0 ∀j

The simulated spatial kinematic-muscular synergies (**W**_**GT**_) simply resulted:

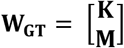

The last step for the creation of the ground truth was to determine the time-varying synergy combination coefficients. This was done employing again the *quadprog* Matlab function, allowing to determine non-negative coefficients ***C*** with minimum norm reproducing, through the synergy matrix **W**_**GT**_ and the moment arm matrix **A**, the time series of joint angle accelerations [26]:

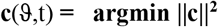

with the constraints **α**(ϑ,t**) = AWc**, *c*_*i*_ ≥ 0 ∀i, and where **α**(ϑ,t) is the vector of angle accelerations for the sample at time t of the reaching movement in direction ϑ. More specifically, we generated **α**(ϑ,t) as follows. First a set of *t* targets in the 4D kinematic space (randomly in the space, Matlab function *rnd*) were selected. Each target vector was then rescaled to Euclidean norm 1, so that the targets lay on 4D-hypersphere of radius 1. For each of the targets, we generated biomimetic joint minimum-jerk motion laws along a straight path in the 4D space [q_i_(t)] connecting the origin to the target. Lastly, we differentiated twice q_i_(t), to achieve **α**(ϑ,t). This procedure can be used for any *n*-dimensional space and *t* target. This allowed us to generate multiple sets of **W**_**GT**_ and **C**_**GT**_.

An example of 4D ground truth **[W_GT_ C_GT_]** with 5 synergies is reported in Fig. 2. Five is the minimum number of synergies (bases) required to ensure complete spanning of the original 4D space [18], [25] and was selected for the simulation.

**Figure 2:**
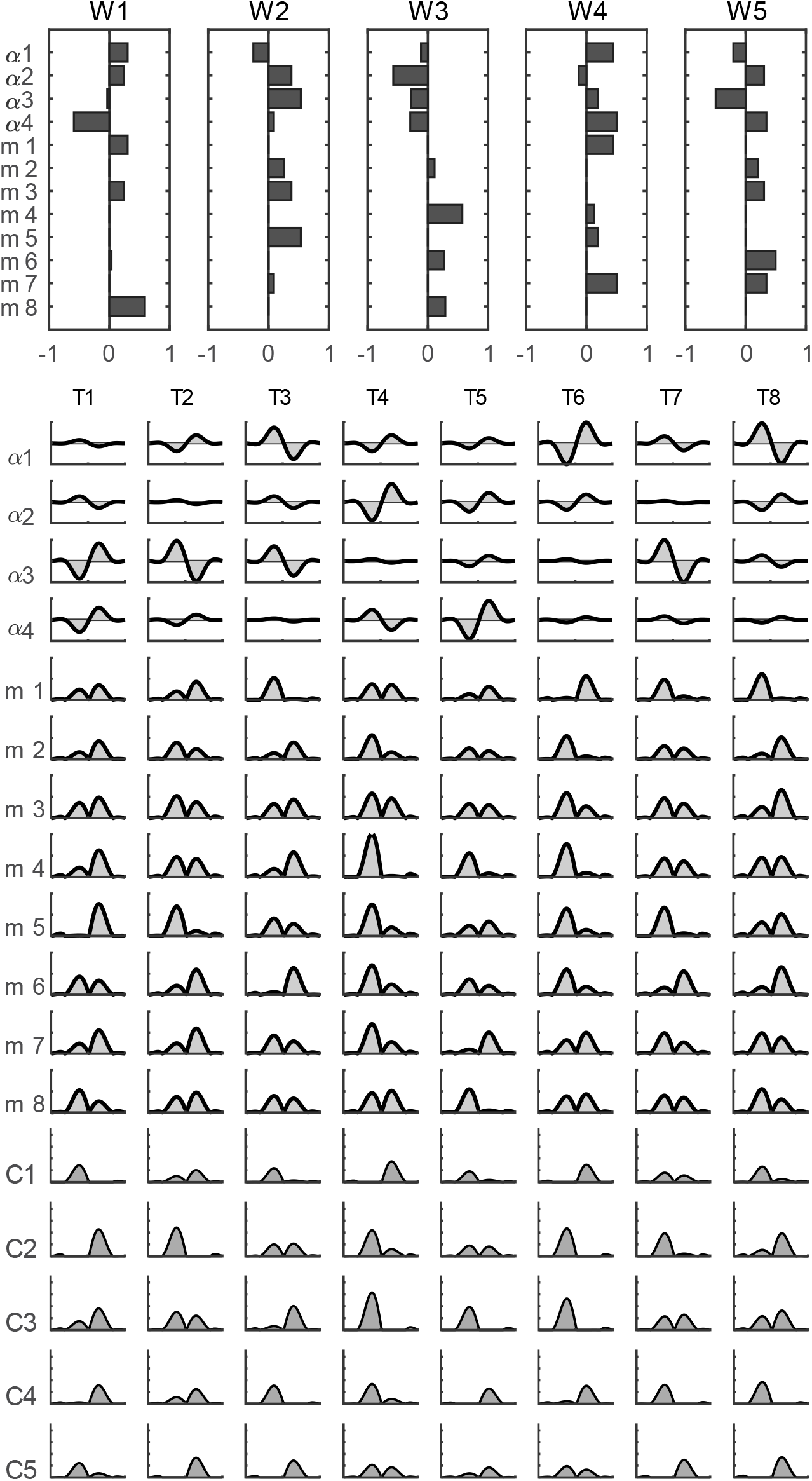
Example of ground truth synergies and combination coefficients. *Top* panel: an example of 5 simulated ground truth kinematic-muscular synergies (**W**_**GT**_); *bottom* panel: simulated time-varying synergy combination coefficients for 8 different targets in the 4D kinematic space (**C**_**GT**_); *middle* panel: simulated data generated by the activation of the synergies modulated by the coefficients (**X = W**_**GT**_ **C**_**GT**_). The dimensionality of the data is 12 (8 muscle activations, 4 joint accelerations).

#### 2.4.2 Cancellations

As mentioned above, when synergy elements are not constrained to be non-negative, there may be cancellations between the elements of different synergies with opposite signs. Once a set of synergies W and temporal coefficients C is given (e.g., **W**_**GT**_ and **C**_**GT**_), they produce a level of cancellation. Thus, we introduced the regularization term in the cost function (with weight λ) to reduce cancellations. We provided a metric (cancellation index, CI) to measure the level of cancellation associated to a W-C set. We defined CI as follows:

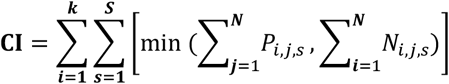

Where *i* is the i^th^ kinematic variable, s is the s^th^ sample of the reconstructed signal, and 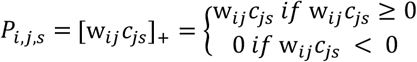 and 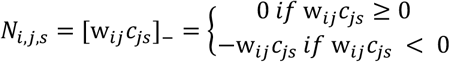 are the positive and negative contributions of synergy *j* on each sample *s* of variable *i* of the reconstructed signal.

#### 2.4.3 Tuning the MMF parameters: λ – *μ* simulation

Before we could perform a full simulation on ground truth data to compare the performance of the MMF and NMFpn algorithms, we ran a preliminary simulation to determine adequate ranges of applicability for the parameters of the algorithm: λ and *μ*. Surfaces depicting relevant outcome parameters were achieved by extracting data from some ground truth sets (n=10) with all possible pairs of λ and *μ* spanning the following values: λ = [0 20 50 100 200] and *μ* = [0.01 0.05 0.1 0.2 0.5]. λ-*μ* surfaces were evaluated considering R^2^, synergy spatial similarity, temporal similarity, and cancellation index to determine suitable values for λ and *μ* to be used in the main simulation.

#### 2.4.4 Extracting synergies from simulated data

Following the λ-*μ* simulation (see Results), we selected the following parameters: *λ*^*s*^ = 50; 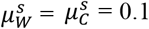. The number of ground truth synergies was set to 5. For each ground truth dataset, we extracted from 1 to 8 synergies. We generated 50 different **[W_GT_ C_GT_]** sets.

Moreover, to verify the robustness of the algorithm to noise, we corrupted the data by additive Gaussian noise **щ** (**X=W_GT_·C_GT_+щ**) according to 5 SNR levels (0 dB, 10 dB, 15 dB, 20 dB 30 dB). Summarizing, we repeated the simulation creating 50 datasets, each one tested with the 5 levels of noise. For each ground truth dataset and level of noise, we performed 20 repetitions of synergy extraction and selected the one with the highest reconstruction **R**^**2**^.

As illustrated in Figure 2, the simulation algorithm is structured with 2 nested loops circling trough a predefined number of datasets and through a set number of noise levels. Additionally, for each dataset and noise level, the algorithm was run multiple times with random initial condition, following the standard procedure for synergy extraction.

The parameters employed were listed in Table 1.

**Table 1:** Parameters for the simulation.

| Parameter | Number | Notes |
| --- | --- | --- |
| Kinematic variables | 4 | Non-constrained |
| Muscle variables | 8 | Non-negative |
| Noise levels | 5 | 30dB, 20dB, 15dB, 10dB, 0 dB |
| Number of datasets | 50 |  |
| Extractions per dataset | 20 |  |
| Total number of extractions | 5000 | $50 \times 20 \times 5$ |

Lastly, the termination condition for the iterations of both algorithms was set according to the decrease in error between two successive iterations and the absolute value of the residual error. When the error between two successive iterations decreased less than a given threshold (τ_1_) amount and the absolute error fraction of the initial error was less that a given second threshold (τ_2_), then the algorithm would stop. In the present work, τ_1_ was set to 10^−3^ and τ_2_ was set to 10^−3^.

### 2.5 Simulation outcome measures and statistical analysis

As a first outcome measure, we tested the capability of the algorithm of identifying the correct number of GT synergies. We employed some of the criteria most commonly adopted in the literature. First, we determined the number of synergies according to two thresholds for the reconstruction **R**^**2**^ (0.80 and 0.90). For each algorithm and noise level, we chose the minimum order needed to reconstruct at least the selected level of reconstruction **R**^**2**^. We also used the linear regression method as in previous studies [18] which identifies the number of synergies as the number at which adding further synergies contributes to the **R**^**2**^ with a constant amount. We performed a linear fitting of the reconstruction **R**^**2**^ curve (Matlab polyfit) from a given order *N* to the maximum (*k + n* variables) and chose the minimum order *N* for which the mean square error of the fit was < 10^−2^, indicating that the “tail” of the curve after the “knee” was essentially straight (as in [18]). We also assessed the performance of the algorithm in recovering the ground truth and reconstructing the simulated data as a function of the amount of noise. As outcome measures, we report the reconstruction **R**^**2**^, the similarity between extracted and GT synergies, and the similarity between extracted and GT synergy combination coefficients. The similarity between the **W**_**E**_ and **C**_**E**_ extracted by the algorithm with the ground truth **W**_**GT**_ and **C**_**GT**_ was computed as:

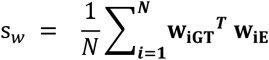

where *i* the i^th^ synergy pair after synergy matching between the two datasets. Synergies were matched by selecting the pair with the highest scalar product (cosine similarity) between all the ***W***_***GT***_ and ***W***_***E***_ pairs and then discarding the matched synergies from the dataset, and repeating iteratively this procedure until no synergy was left (all synergies were matched). Since **w**_**iGT**_ and **w**_**iE**_ have unit norm, -1≤ **s**_**w**_ ≤1. Similarly, the similarity between synergy combination coefficient was computed as the mean correlation coefficient,

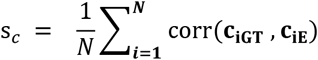

computed on the temporal coefficients of the matched pairs. This computation was repeated for each **W**_**E**_ and **C**_**E**_ extracted from each ground truth set and noise level. Then, for each noise level, we generated a distribution of the best similarities achieved across ground truth sets.

To assess the difference in performance between the MMF and the NMFpn, we used a Kruskal Wallis test, on the results achieved with both algorithms on **W, C**, and reconstruction **R**^**2**^. Tests were repeated for each noise level. Post hoc tests were conducted with Matlab *multcompare*.

### 2.6 Test on upper-limb muscle-kinematic real data

We tested the performance of the novel algorithm in a real scenario involving the analysis of upper-limb reaching movements by extracting muscular-kinematic synergies from muscle activation patterns and joint angle acceleration waveforms, using a similar methodology to the dataset collected in a previous study [26].

#### Participant

We collected both muscular and kinematic variables collected from a participant executing point-to-point upper-limb movements. The participant was a 35-year-old male, 81 kg of weight, and 1.80 m of height. The data are within a study reviewed and approved by the CNR Ethical Committee (Rome, Italy, protocol number 0044338/2018). The participant provided written informed consent to participate in this study.

#### Experimental Set up

The tests were performed in the motion acquisition laboratory of the Italian Council of National Research (CNR) in Italy, Lecco. The laboratory equipment included a Vicon Vero system, with 10 infrared cameras and a set of reflective markers for motion tracking, and a set of EMG probes (Cometa wireless, 16 channels). In this experiment, a 34 markers set was used (25 for the Vicon upper limb model was used, 9 for the target) and 14 EMG probes were used. A board with 8 targets arranged on a circle (60 cm diameter) at 45° angular distance in points of interest. The target board was placed in the sagittal plane with respect to the experimental subject, similarly to a previous study [3]. Targets were labelled, clockwise: FW-UP (forward-up), FW (forward), FW-DW (forward-down), DW (down), BK-DW (backward-down), BK (backward), BK-UP (backward-up), UP (up). An additional target at the centre of the board (labelled O) indicated the start position. The target board and the participant were placed in the acquisition volume of the Vicon system. A schematic illustration of the setup is portrayed in Fig. 3. Movements were performed with the dominant upper limb (right).

**Figure 3:**
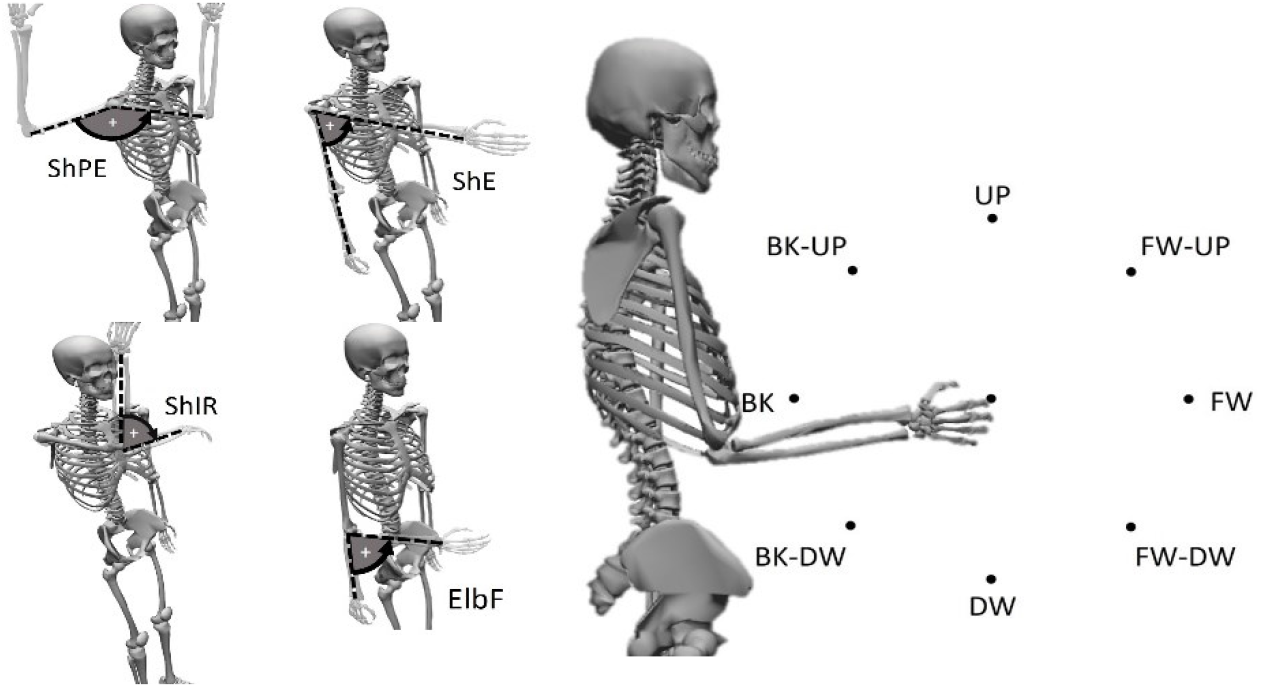
Joint angles and targets. *Left*: The following joint angles were considered: Shoulder Plane of Elevation (ShPE: + for medial rotations, - for lateral rotation); Shoulder Elevation (ShE): + when elevating the limb, - when going in the direction of gravity); Shoulder Internal Rotation (ShI: + internal rotation, - external rotation); Elbow Flexion (ElF: + when flexing, - when extending). *Right*: Targets for center-out movements, clockwise: FW-UP (forward-up), FW (forward), FW-DW (forward-down), DW (down), BK-DW (backward-down), BK (backward), BK-UP (backward-up), UP (up) and 1 target in the centre (labelled O).

#### Experimental protocol

Point-to-point movements involve multi-joint coordination and are used in daily life activities. They are also often used by clinicians to test motor capabilities of patients. In our experimental protocol, the subject started from a predefined resting position, with the arm leaning along the body and with the elbow flexed and the forearm horizontal, the hand in neutral position, with the forearm neither pronated nor supinated. Starting from the 0 target, the subject had to reach toward the first target (FW-UP) of the board, and then return to the starting position (0). Then, the subject continued the center-out/out-center motion pattern toward all targets, clockwise, until he reached the UP target, and backward to target 0 to conclude a trial.

#### Data acquisition

Before each recording session, the subject was instrumented with markers in accordance with the standard upper limb model designed for the Vicon system. Five markers were placed on the trunk, one on each shoulder, three on each upper arm, two on each elbow and four on each forearm and wrist [27], for a total of 25 markers. Kinematic sampling frequency was 100 Hz. In order to collect muscular data, a set of 14 EMG probes were positioned on the torso and upper-limb of the subject to sample EMG signals at 1000Hz. The tracked muscles were the following: Latissimus Dorsi (LatD), Teres Major (Ter M), Lower Upper Trapezius (UT), Infraspinatus (Inf S), Deltoid Anterior (DA), Deltoid Middle (DM), Deltoid Posterior (DP), Pectoralis Clavicular Head (Pect), Triceps Long Head (TriL), Triceps Lateral Head (TriLa), Biceps Long Head (BicL), Biceps Short Head (BicS), Pronator Teres (PR), Brachioradialis (Brd).

The upper limb kinematic model implemented in the Nexus 2.1 software allowed the reconstruction of angular coordinates of the glenohumeral joint centre (shoulder centre of rotation), humeroulnar joint centre (elbow centre of rotation) and radiocarpal joint centre (wrist joint centre). Using these data, we selected a set of 4 degrees of freedom describing the motion of proximal upper limb joints with the following conventions: q1 = Shoulder Plane of Elevation (ShPE: + for forward rotations, - for backward rotation); q2 = Shoulder Elevation (ShE):+ when elevating the arm, - when depressing the arm); q3 = Shoulder Internal Rotation (ShIR: + internal rotation, - external rotation); q4 = Elbow Flexion (ElbF: + when flexing, - when extending). For clarity, they are reported in Fig. 3. All further analyses were conducted in MATLAB (MathWorks, Natick, USA).

#### Data analysis

Let a trial be the concatenated movements along all the cardinal directions of the board. Both kinematic and EMG data recorded in each trial were pre-processed by filtering, segmentation, and normalization. During the acquisition, the Vicon system (kinematics) and EMG system (Cometa) were synchronized via hardware from the manufacturer (the same trigger started and stopped simultaneous acquisitions with both signals). All EMG data were delayed of 50 ms to account for an estimated electromechanical delay between the signals [28].

Marker positions were filtered with a low-pass Butterworth filter with a 15 Hz cut-off frequency, 5^th^ order. Both kinematic and EMG data were then segmented in phases (a center-out movement from target 0 to a peripheral target, an out-center movement from a peripheral target 0) based on the tangential velocity profile of the end effector (wrist joint centre). The beginning of each phase was defined as 200 ms before the time at which the velocity reached 5% of the peak velocity; the end of each phase as 200 ms after the time at which the velocity dropped to 5% of the peak velocity. All kinematic data were derived twice to achieve joint angular accelerations. Lastly, each angular acceleration was normalized with respect to the maximum of its absolute value. This allowed to rescale accelerations in a range between -1 and 1. Only center-out movements were included in the analysis.

EMG signals were filtered to obtain smooth envelopes and to remove any potential motion artefacts. The applied pre-processing steps were the following: the signal was filtered with a third order high-pass Butterworth filter with 3 Hz cut-off frequency, full-wave rectified, and filtered with a third order low-pass Butterworth filter with 15 Hz cut-off frequency. After filtering, the EMG data was segmented in phases using the same events obtained from the kinematics. Additionally, in each phase, the tonic component of the EMG signal was removed by subtracting a linear ramp according to previously employed models [29]. Then, each EMG envelope was normalized to its maximum absolute value and multiplied by This choice was made to make EMG values range [0, 2] comparable with the one of the kinematic variables [-1, +1]. In this way, each EMG waveform was scaled, achieving the normalized EMG envelopes. To account for negative waveforms achieved after tonic subtraction (minimal in magnitude), negative EMG data were clipped to 0 as in previous works [18], [26]. Lastly, the kinematic and EMG signal in each phase were re-sampled at 100 samples and averaged over 10 repetitions for each target.

The filtered, segmented, resampled, and averaged kinematic data (angular accelerations of 4 upper-limb degrees of freedom) and EMG data (14 channels) were grouped in an aggregated muscle-kinematic matrix (MKM) before synergy extraction. The MKM had dimensions 18 × 800, where 18 is the sum of 4 selected kinematic and 14 muscular variables, and 8 directions were considered, each one having 100 samples.

Lastly, the MMF algorithm was used for extracting synergies with the setting shown previously: *λ* = 50, *μ* = 0.1, number of extracted synergies ranging from 1 to 14.

## 3. Results

### 3.1 Selection of algorithm parameters

To select adequate parameters of the algorithms, i.e., learning rates (μ) and weight for the regularization cost (λ), we performed multiple simulations with data generated by combinations of know synergies and combination coefficients (ground truth). We tested the effect of varying μ and λ on the quality of the reconstruction (R^2^), amount of cancellation, similarity of extracted spatial synergies with ground truth synergies, and similarity of extracted synergy combination coefficients with ground truth coefficients (Fig. 4). As expected, R^2^ was maximal for λ = 0; however, increasing λ allows to reduce the amount of cancellation leading to more physiologically plausible solutions. This can be achieved in a large range of λ values with minimal effect on R^2^. Consequently, using a reasonable λ is advantageous for the physiological interpretation of the extracted synergies. Although spatial and temporal synergies similarity slightly decreased when increasing λ they were almost always very high. Only when λ or μ were at their maximum tested value the spatial and temporal similarity decreased consistently. We concluded that a reasonable choice for our simulation was to select λ = 50 and μ = 0.1. These values allowed to reduce cancellations (toward level similar to the ground truth) without altering reconstruction accuracy.

**Figure 4:**
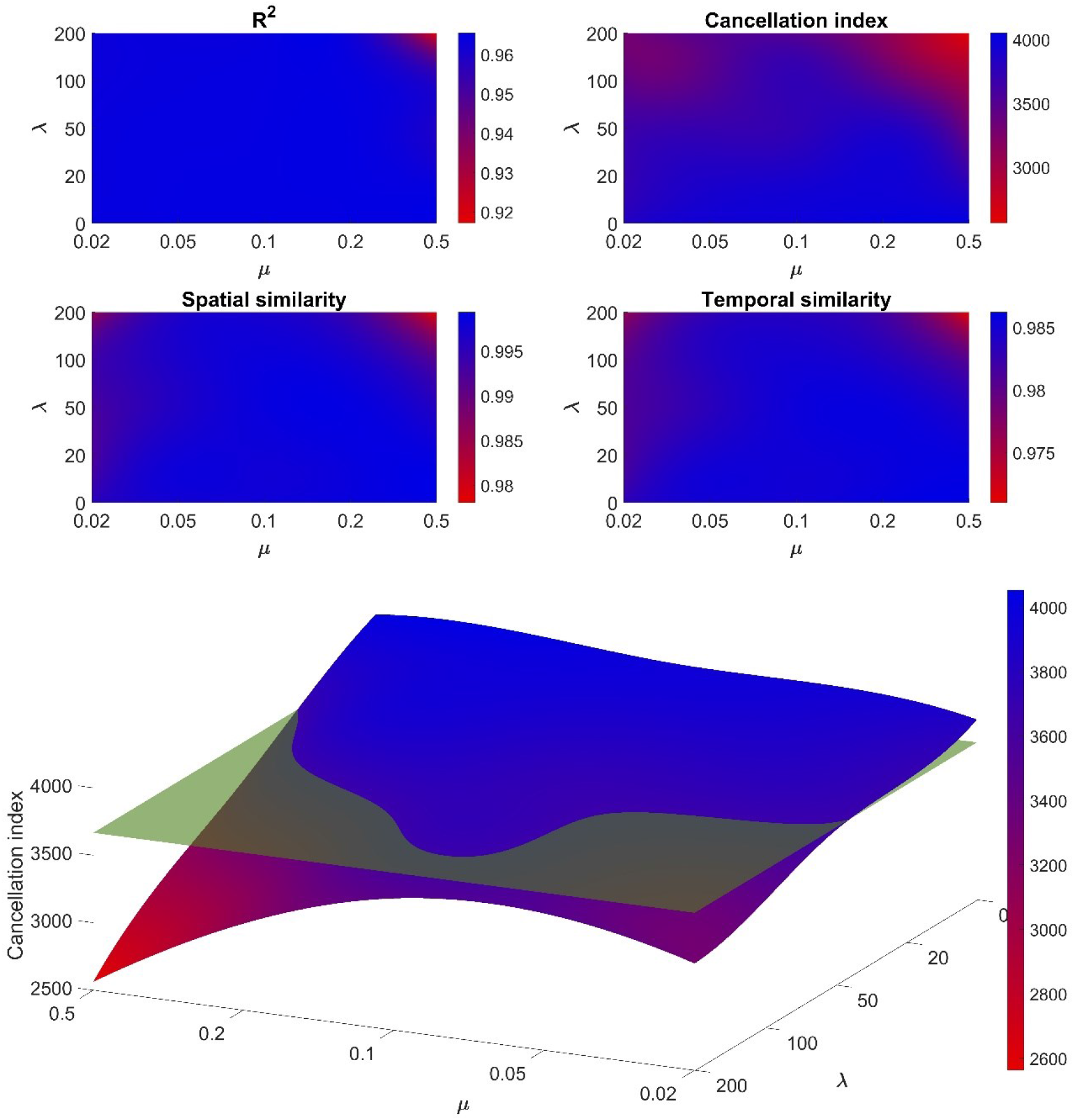
Dependence of the reconstruction R^2^, cancellation, spatial synergies similarity, temporal synergies similarity on algorithm parameters (*λ* and *μ*). The plots illustrate the R^2^ values achieved when varying *λ* and *μ* (*top left* panel) and the Cancellation Index (*top right* panel). Some regions, corresponding especially to high values of *λ*, can reduce cancellations but this effect is achieved with a reduction of spatial (*bottom left*) and temporal (*bottom right*) similarity of the extracted synergies. In the *lower* panel, a 3D surface reproduced the data of panel b together with the average level of cancellation of the ground truth data. We selected as values for the MMF-NMFpn simulations the following parameters: λ=50 and μ=0.1.

### 3.2 Validation with simulated data

In Figure 5, we report the distributions of the R^2^ values for the reconstruction of the data generated with the combination of 5 ground truth synergies (each one including 4 kinematic variables and 8 muscular variables) when varying the number of extracted synergies.

**Figure 5:**
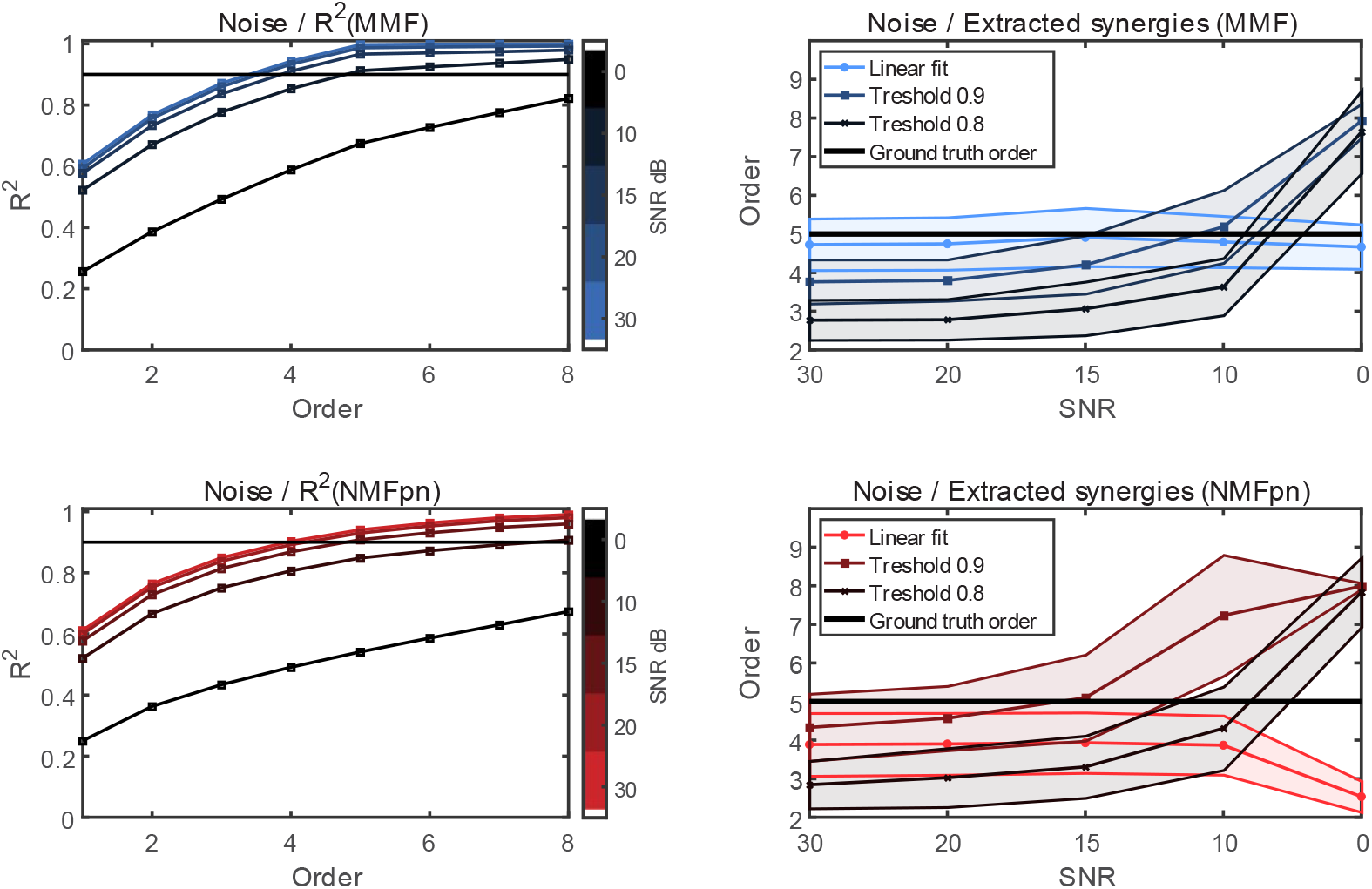
Effect of noise on R^2^ curves and selection of number of synergies. The ground truth data is based on a set of 5-dimensional set of bases (kinematic-muscular synergies). The number of extracted synergies is determined for both MMF (*top left*) and NMFpn (*top right*) using two R^2^ threshold criteria and a linear fit criterion (number of extracted synergies is the minimum order for which the R^2^ curve can be linearly interpolated with RMS below 10^−2^). With MMF and the linear fit criterion it was possible to identify the correct number of ground truth synergies (*bottom left*) in all noise except, while NMFpn had lower performance (*bottom right*).

The number of extracted synergies was determined for both the MMF and NMFpn according to three different criteria: (1) using a R^2^ threshold of 0.8, (2) using a R^2^ threshold of 0.9, (3) using a criterion based on the flattening of the R^2^ curve estimated by linear regression (i.e., the number of synergies selected is the minimum number for which the R^2^ curve from that number to the maximum number can be linearly fitted with a RMS error below 10^−2^). Results are reported for all the 5 levels of noise tested. We note that with MMF, using the linear regression criterion, we were able to select almost in all cases the correct number of ground truth synergies, even with large amount of noise corruption (i.e. up to SNR = 0 dB).

In Fig. 6, we report the main results of the simulation for a number of extracted synergies corresponding to the ground truth (5 synergies). In Fig. 6 (*top left*), we report the reconstruction R^2^ on simulated data comparing the MMF and the NMFpn. Interestingly, for the condition with lowest noise (SNR = 30dB), the reconstruction achieved with the MMF algorithm was almost perfect (R^2^ = 1). The Kruskal-Wallis test revealed a significant effect of noise on the reconstruction R^2^ for both MMF and NMFpn (p<0.001). Remarkably, MMF performed always significantly better than the NMFpm for all SNR levels (p < 0.001). However, we note that even at low SNR (10 dB), the MMF algorithm performed well (R^2^ > 0.90).

**Figure 6:**
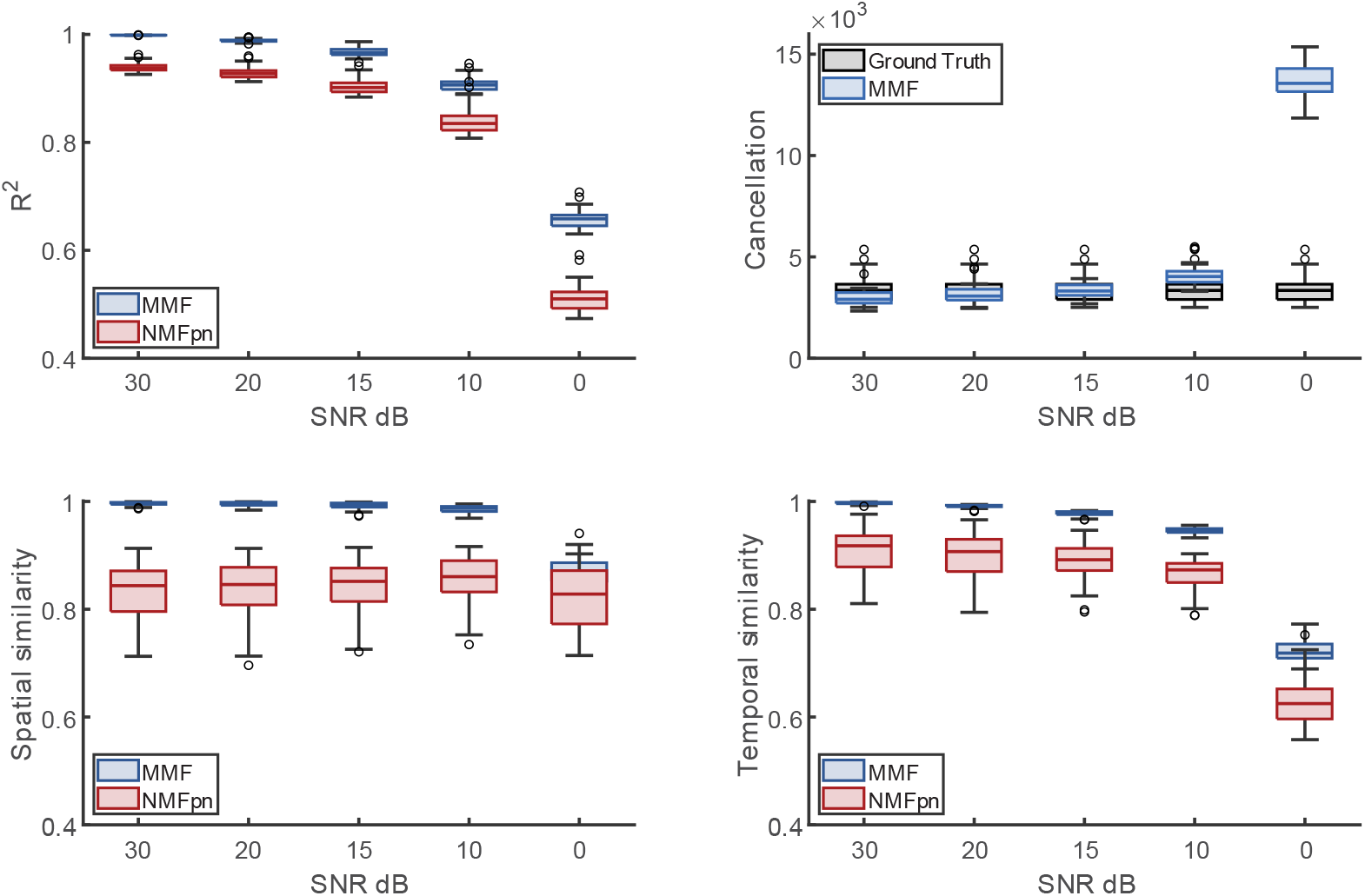
Distribution of reconstruction R^2^, cancellation index, spatial synergy similarity and temporal synergy similarity for different noise levels. Boxplots for the distributions of R^2^ (*top left*), cancellation index (*top right*), spatial synergy similarity (*bottom left*), and temporal coefficients similarity (*bottom right*) achieved by the MMF (*blue*) and NMFpn (*red*) algorithms with different noise levels. Data for 4 kinematic DoF and 8 muscles are simulated as combinations of 5 ground truth kinematic-muscular synergies.

In Figure 6 (*top right*), we report the distributions of the similarity between the extracted synergies and ground truth spatial synergies (**s**_**w**_). Interestingly, for SNR = 30dB, spatial synergies similarity (**s**_**w**_) with MMF was almost perfect. Increasing the noise, the similarity remained high even reducing SNR to 10 dB. Similarly, **s**_**w**_ achieved with NMFpn did not depend on noise (except for the “extreme” case with SNR = 0 dB) but is always below 0.90. MMF performed always better than NMFpn in a statistically significant way for all SNR levels (p<0.001). In Figure 6 (*bottom left*), we report the similarity between extracted and ground truth synergy temporal coefficients (**s**_**c**_). Interestingly, for SNR = 30dB, **s**_**c**_ achieved with MMF was almost perfect. Increasing the noise, the similarity slightly decreased but remained well above 0.90 even when reducing SNR to 10dB. Similarly, with NMFpn, **c**_**w**_ was only slightly dependent on noise but was below the values achieved with MMF in a statistically significant way for all SNR levels (p<0.001). In Figure 6 (*bottom right*), we report the cancellation index. In “medium” noise conditions (SNR = 15dB) the MMF algorithm can reproduce quite accurately the level of cancellation found in the ground truth. From this graph, it can be deduced that when the SNR decreased, an increasing value for λ was needed to reproduce the cancellation of the ground truth. λ =50 worked well for our conditions at 15dB and 20dB, while it was too low for 10dB, extremely low for 0dB, and too high for 30dB.

In Figure 7, we illustrate an example of the performances of the MMF and NMFpn algorithms on simulated data (SNR = 15 dB). The R^2^ curve for the MMF algorithm shows a clear slope change allowing to identify 5 synergies as in the ground truth. Accuracy on the reconstructed spatial synergies and temporal coefficients is also much higher with the MMF. Data reconstruction is almost perfect with MMF, while NMFpn cannot reconstruct well the kinematics, slightly altering the original acceleration waveforms.

**Figure 7:**
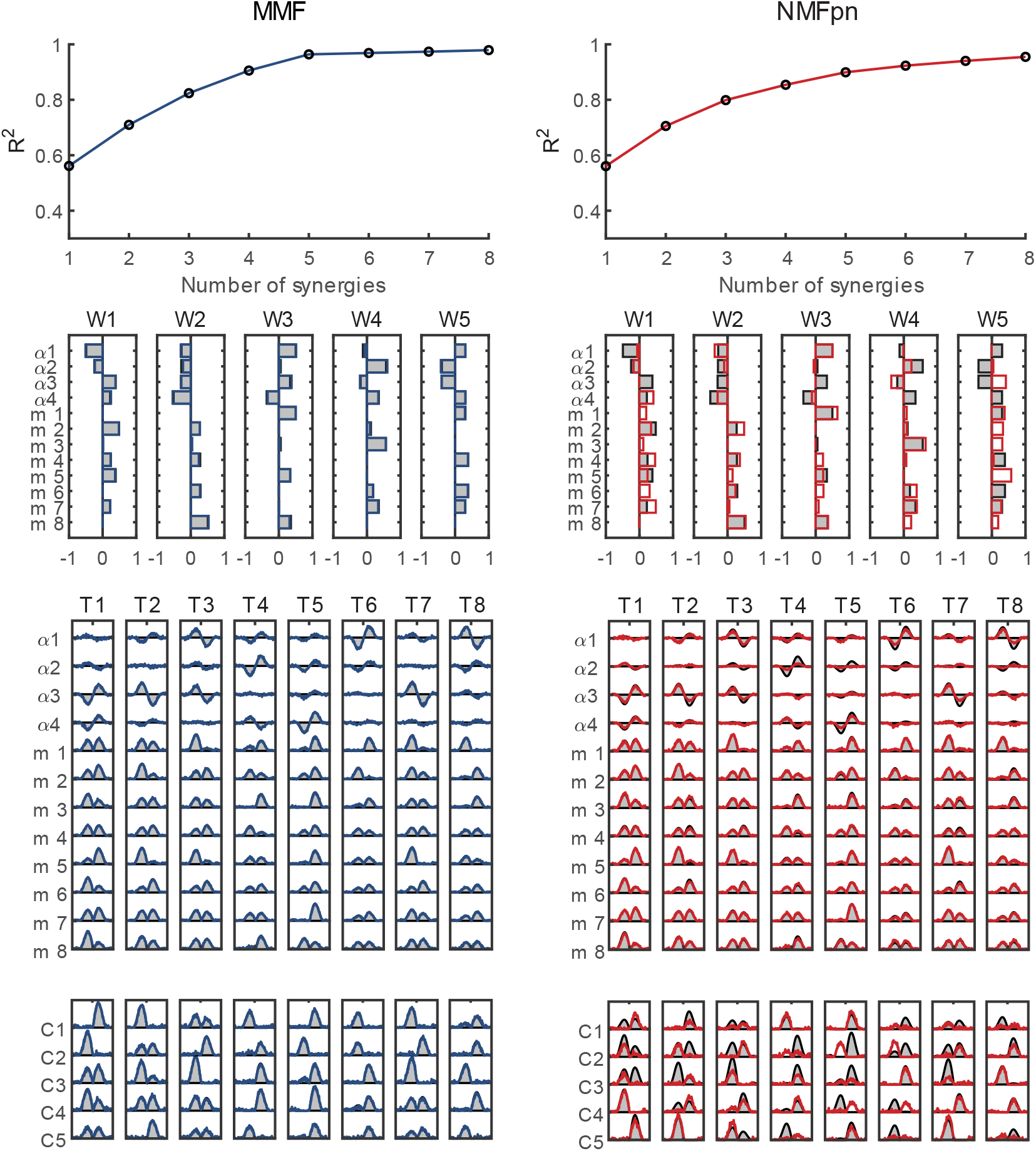
R^2^, extracted synergies and temporal coefficients for one extraction from simulated data: MMF-NMFpn comparison. An example of the R^2^, spatial synergies, reconstructed signal, and temporal coefficients extracted with the MMF algorithm (*left*; R^2^: *top, blue line*; synergies: *blue bars*; temporal coefficients: *bottom, blue lines*; reconstructed signal: *blue line*s) and with NMFpn (*right*, R^2^: *top, red line*; synergies: *red bars*; temporal coefficients: *red lines*; reconstructed signal: *bottom, red lines*), both compared to the original ground truth (synergies: *black thick bars*; temporal coefficients: *grey-coloured areas*; reconstructed signal: *grey-coloured areas*) for one representative dataset with SNR = 15dB, λ = 50.

### 3.3 Application to real data

The MMF algorithm was used to extract kinematic-muscular synergies from point-to-point center-out movements towards 8 different targets on the sagittal plane (Fig. 8).

**Figure 8:**
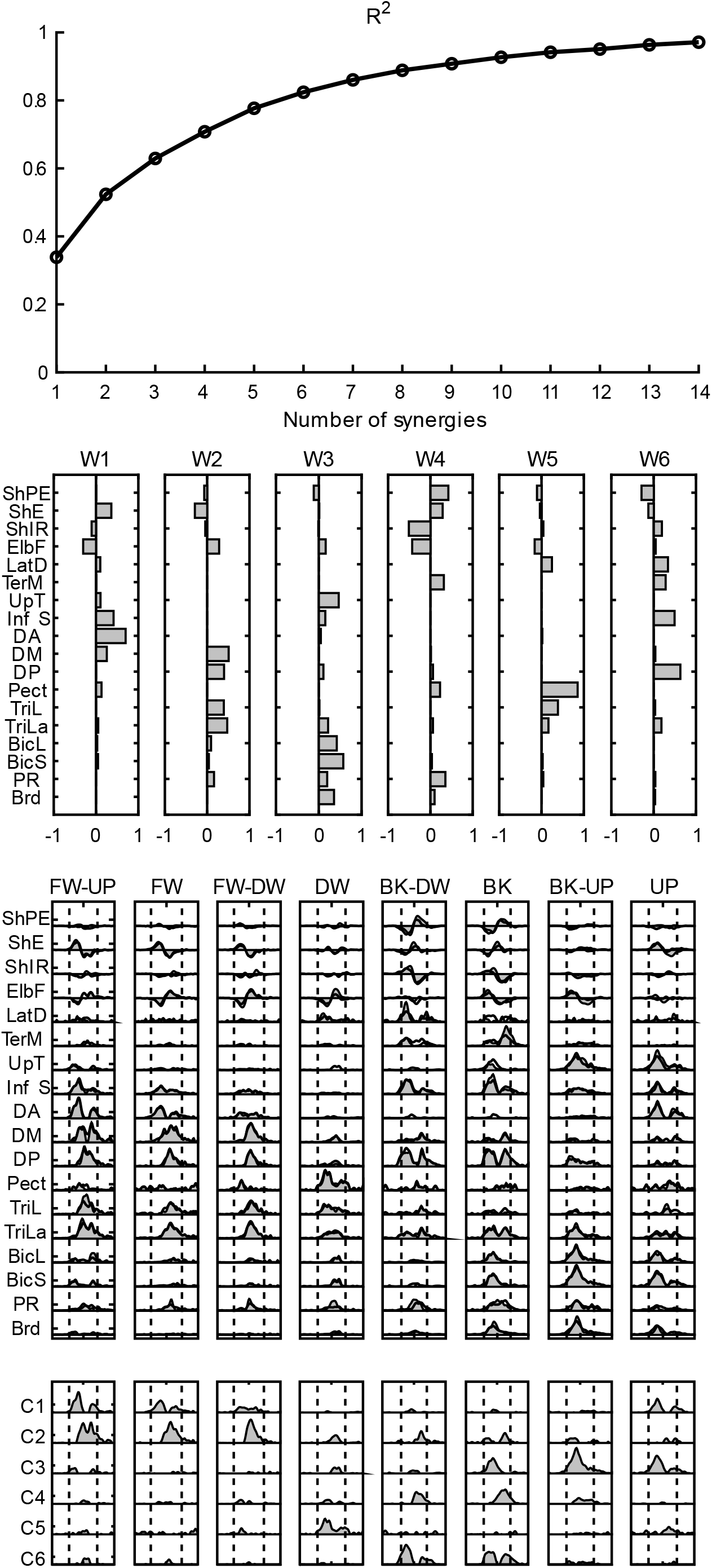
Application of MMF to experimental data. Data collected during reaching movements to 8 targets (*centre-out movements*) arranged in the sagittal plane were used to extract kinematic-muscular synergies and combination coefficients. In the *top* panel, the curve of the reconstruction R^2^ value as a function of the number of extracted synergies is shown. In the *top-middle* panel, the extracted kinematic-muscular synergies are portrayed. In the *middle* panel, the original data (grey areas) and reconstructed data (red line) are portrayed. In the *bottom* panel, the coefficients used to modulate synergies are reported. Notation: ShPE = Shoulder Plane of Elevation; ShE = Shoulder Elevation; ShIE = Shoulder Internal Rotation; ElbF = Elbow Flexion (see Methods). Movement onset and offset are shown with vertical lines. Movement onset and offset are shown with vertical lines.

Six synergies were selected based on the reconstruction R^2^ curve using the linear regression criterion [18]. The synergy number is slightly higher than the one used for the simulation, but the number of muscles included (the most relevant muscles involved in sagittal point-to-point movements) is larger (14 here vs. 8 in the simulation). Since all movements are point-to-point, we expected to find biphasic kinematic-muscular links, related to the acceleration or the deceleration phase typical of this class of movements [18]. Moreover, for each direction of motion, we expected that at least two synergies would act for achieving acceleration and deceleration patterns that cannot be obtained with a single synergy (even if, in principle, the same synergy can participate to both the acceleration and deceleration phase). This was indeed observed in our data. The temporal coefficients (C) are modulated so that at least two kinematic-muscle synergies are properly timed and coordinated to reconstruct the original patterns for all the directions of motion. It is for example the case of W1 and W2, that are “antagonistic” synergies. W1 acts in the agonist phase to elevate the limb and extending the elbow toward the forward-down (FW-DW), forward (FW) and forward-up (FW-UP) directions, while W2 intervenes to decelerate shoulder elevation and counteracts elbow extension (by flexing the elbow). Moreover, most of the synergies show quite clear modulation across end-point movement directions with distinguishable peaks in their directional tuning, accounting for each acceleration and deceleration phase in a plausible way. Temporal coefficients are mostly either biphasic or monophasic, and related to the acceleration or the deceleration phase. We also noted that not all synergies can be easily provided with a clear physiological interpretation. It is the case of W4, that shows a dominance of kinematic patterns in the backward (BK) and backward-down (BK-DW) directions.

## 4. Discussion

### Summary of the main findings

We introduced a novel algorithm to perform a matrix factorization of data that contains a mixture of both non-negative and unconstrained variables and, thus, can be employed to extract kinematic-muscular synergies. The MMF algorithm overcomes some of the limitations of previous attempts of relating synergistic approaches applied to EMG and kinematics, usually analysed separately. We first identified proper ranges for the parameters of the algorithm. Then, we validated the algorithm on simulated data generated by the combination of known synergies and combination coefficients. We generated such ground truth using a simple linear mapping of muscle activation to kinematic variables. We found that the algorithm accurately recovered the ground truth from noiseless data. When noise was added, the MMF algorithm correctly identified the correct number of synergies in most cases (using as criterion the flattening of R^2^ curve) in most cases. The similarity of the extracted synergies and combination coefficients with the ground truth synergies and coefficients decreased with increasing noise but was always higher than the similarity achieved with the NMFpn approach. We then tested the algorithm on real data and we found that the extracted kinematic-muscular synergies capture meaningful relationship between shoulder and arm muscles and hand accelerations.

### Simulation results

The results of our simulation fully validated the MMF algorithm. With noiseless data, we were able to achieve a reconstruction R^2^ stably over 0.99, meaning that the algorithm was able to reconstruct almost perfectly the original kinematic-EMG signals. While adding noise significantly decreased the reconstruction R^2^, the reconstruction quality was always very high and was always higher with MMF rather than with NMFpn. When assessing similarity of spatial synergies and temporal components, results were equally supportive of the validity of the algorithm. In fact, when no noise was added to the data, we could reconstruct the original **W**_**GT**_ and **C**_**GT**_ almost perfectly [s_w_>0.99, s_c_>0.99]. When increasing noise, as expected, there was a slight drop of the similarity values. The drop is in any case limited and reconstruction metrics were always reliably high, even in presence of noise. Even with these metrics, we also noted that the performance of the MMF algorithm was always higher than NMFpn. We were also able to reconstruct effectively the original number of synergies by using the linear regression method which we suggest to couple with the employment of the MMF algorithm.

### Selection of the parameters of the algorithm: λ and μ

In this study, we selected adequate values for λ *and* μ according to tests on simulated data. It is indeed of interest to comment on these parameters. Increasing μ results in a higher gradient step, which may lead to faster convergence but may miss more frequently the global minimum. On the contrary, decreasing μ may require too many iterations to converge to the solution. λ played instead a crucial role in reducing cancellations and forcing sparseness on the solution. In fact, when simulating data and extracting synergies with λ = 0 we observed large cancellations between kinematic components of different synergies with opposite signs, recruited simultaneously. In this condition, the reconstructed signal may not be perfectly decomposed into the original ground truth, and this is reflected in our findings. We note that this condition of ambiguity is different from NMF where contributions from different synergies can only add up, but they cannot be cancelled. Cancellations take place when synergy components have different signs, and when their temporal coefficients are active with overlapped timings.

One of our findings is that the algorithm without regularizing term λ can in general (and often does) find solutions associated with high levels of cancellation. Of course, while this is a minimum for a cost function, these solutions cannot be accepted since they likely do not reflect the organization at kinematic-muscular synergy level. In fact, slightly increasing λ has the effect of reducing the amount of cancellation at the cost of a slightly decrease in the reconstruction R^2^, which is perfectly acceptable within some ranges quantified in the paper and depending on analysed dataset.

### Comparison with previous approaches

Torres-Oviedo and collaborators [22] used NMF to extract functional muscle synergies from combined muscular (non-negative) data and force (unconstrained) data. While functional muscle synergies extracted from separate positive and negative waveforms (NMFpm) robustly captured muscle activations and the active force vectors produced during postural responses under several biomechanically distinct conditions, it is not clear whether they represent an optimal set of synergies. In fact, the separation of the endpoint forces into positive and negative components along Cartesian axes may lead to an incorrect estimation of the synergy set dimensionality. When mapping a set of force vectors whose non-negative combinations span the entire force space into vectors of the positive/negative component space there is no guarantee that they will still span the entire force space. For example, considering for simplicity two dimensional vectors ([x y]), non-negative combinations of the vectors {[1 0], [0 1], [-1 -1]} span the entire space but the non-negative combinations of the corresponding four dimensional positive/negative component vectors ([x+ x-y+ y-]) {[1 0 0 0], [0 0 1 0], [0 1 0 1]} do not span the space. In fact, the original vector [1 -1], corresponding to [1 0 0 1] in the space with separate positive/negative components, cannot be generated by any non-negative combination of the three positive/negative component vectors. Thus, at least four different positive/negative component vectors are necessary for spanning a two-dimensional space with non-negative combinations while three vectors are sufficient in the original space. Moreover, the region of original space not spanned by the positive/negative components depends on the choice of the coordinates system.

The limitations of the NMFpn approach were clearly highlighted in our simulation. Indeed, even for noiseless data, NMFpn could not accurately recover the ground truth. When noise was added, the similarity of the synergies and combination coefficients extracted by NMFpn with the ground truth was always lower than the similarity achieved by MMF. Moreover, NMFpm failed to correctly identify the number of ground truth synergies generating the data.

### Physiological interpretation of real data

To demonstrate the applicability of the MMF algorithm to real data, we considered point-to-point multi-directional reaching in the sagittal plane, frequently employed in generic motion-analysis scenarios [26], rehabilitation, and test of devices [30]. We are aware that we are presenting just an example of applicability and that reliable testing of the novel algorithm should be provided on more structured datasets [31], [32]. Nonetheless, in the reported example, despite the limited directional variability, most of the extracted synergies show meaningful kinematic-muscular relationships, reflect the structure of the original input waveforms, and capture well the biphasic nature of point-to-point movements. As the acceleration profiles present a biphasic pattern, with acceleration and deceleration phases [18], [33], non-negative recruitment of kinematic-muscular synergies requires a sequencing of at least two synergies with acceleration components of opposite signs. This can be seen in all the considered kinematic degrees of freedom with different modulation depending on directions. For example, in the Forward, Forward-Up and Up directions, W1, which has a positive loading of shoulder elevation associated with the activation of the anterior deltoid muscle, is recruited first (e.g. C1 has an earlier activation peak) and W2, which has a negative loading of shoulder elevation associated with the activation of middle and posterior deltoid, is recruited later (e.g. C2 has a later activation peak). However, not all synergies have a clear and straightforward physiological interpretation, probably due to the approximations introduced in modelling the kinematic-EMG relationship with a linear model. We observed some synergies in which kinematic and muscular activation are not equally balanced, especially in the antagonist phases. For example, the structure of W4, which loads heavily kinematic variables and only a few muscles, may be affected by the need to capture the biphasic kinematic patterns in two directions (BK and BK-DW). This is not surprising considering that we are approximating the EMG-acceleration relationship with a linear model that does not capture the complex multi-joint dynamics. For this reason, probably our upper-limb scenario is very challenging for the application of the kinematic-muscle synergies. However, it was in our opinion valuable to show some of the potential of the algorithm in a typical scenario that has been often analysed only in the framework of muscle synergies.

Overall, despite some limitations and open issues that require further investigation, these results are promising when considering real applications of the MMF to kinematic-muscular synergies. Even if our approach is merely phenomenological and cannot be compared to a rigorous biomechanical modelling, the versatility of our model may be equally effective in representing relationships between EMGs and joint angular velocities, or EMGs and articular ranges of motion, or pushing this concept to the limit, also to other domains. All these aspects are worth being investigated in future applications of the algorithm.

Finally, the role of some pre-processing choices needs to be highlighted. First, a fixed electromechanical delay was used but it may vary across muscles and it may be necessary to adjusted it differently for each muscle. Secondly, the choice of normalization across domains may affect the results. In this preliminary application, we normalized each muscular and kinematic variable by its maximum value, similarly to a recent experiment [26]; however, which normalization method is more adequate to detect links between domains should be analysed in more detail. Also, the choice of relative scaling between muscular and kinematic data should be assessed. In this study, we decided to preserve maximum ranges but this choice should be discussed and tested in more detail. Still, it is always possible to reconstruct original waveforms by rescaling after extraction. The best and most suitable options for applicability in real scenarios will be matter for further studies.

### Other applications

The MMF algorithm has the potential to improve the currently available methods for extracting muscle synergies, and especially, to extend them to multi-domain scenarios. In particular, the method was designed to capture the relationships between the EMG and the joint accelerations, extending the concept of muscle synergies to muscle-kinematic synergies. However, this novel algorithm is not necessarily limited to this type of analysis. It can be applied to a variety of multivariate problems. Virtually, the algorithm could be used to extend these findings to describe the EMG/Torque [18] and EMG/Force [23] relationships, giving a description of the dynamic of the system, or linking the EMG to muscle forces. This approach would prevent the approximation of kinematic-muscle synergies that neglect multi-link dynamics (in the presented approach, EMGs are associated to accelerations rather than to torques). The scenarios for application are multiple and might also include muscle synergies after the subtraction of the tonic components (including negative phasic contributions as in [3], and [26]). Thus, we foresee possible applications related to the evaluation of negative EMG phasic components (extending the concepts already considered in [3] and [22]). For muscle synergies, this allows phasic components to be negative [3] while in recent studies, they were constrained (clipped) to 0 to be compatible with NMF [26].

## 5. Conclusions

We introduced a novel algorithm for extracting kinematic-muscular synergies that allows to remove the constrain of non-negativity typical of previous approaches. We described in the detail the algorithm and provided comprehensive assessment of its performances and range of applications. We believe that this contribution will help expanding the use of synergistic approaches to motor control in various fields of research.

## Acknowledgments

Authors wish to thank Cristina Brambilla per her support in laboratory recordings. data availability statement: data for this paper are mainly simulated. Laboratory data are available from authors upon reasonable request.

## Author contributions

*CRediT roles:* Conceptualization: AdA, AS; Data curation RM, AS, AdA; Formal analysis: RM, AS, AdA; Funding acquisition: AS; Investigation: AS, RM, AdA; Methodology: AdA, AS; Project administration: AS, AdA; Resources: AS; Software: RM, AdA, AS; Supervision: AS, AdA; Validation: AS, RM, AdA; Visualization: AS, RM, AdA; Roles/Writing AS, AdA, RM; Writing - review & editing AS, AdA, RM.

